# SAMPL: The Spreading Activation and Memory PLasticity Model

**DOI:** 10.1101/778563

**Authors:** Beau Sievers, Ida Momennejad

## Abstract

We present the Spreading Activation and Memory PLasticity Model (SAMPL), a computational model of how memory retrieval changes memories. SAMPL restructures memory networks as a function of spreading activation and plasticity. Memory networks are represented as graphs of items in which edge weights capture the strength of association between items. When an item is retrieved, activation spreads across nodes depending on edge weights and the strength of initial activation. A non-monotonic plasticity rule, in turn, updates edge weights following activation. SAMPL simulates human memory phenomena across a number of experiments including retrieval induced forgetting, context-based memory enhancement, and memory synchronization in conversational networks. Our results have implications for theorizing memory disorders such as PTSD and designing computationally assisted conversational therapy.

## Introduction

Memory is malleable. Remembering something directly, like an item, a person, or an event, enhances our memory of it. Memory is organized in relational structures^5^. When we remember something (a friend’s name), we automatically retrieve related memories. Some of these memories are closely related (what the friend looks like) and some are further away (a mutual friend). This “spread of activation”^1^ across a network of memories can change memory for related items^1–4^. However, while some memory studies have shown that memory for related items is enhanced^1^, other studies have shown that remembering something can weaken memory for items that are related to it^1–4^. This apparent contradiction poses challenges to models of memory retrieval. One challenge is to offer unifying memory principles that explain how and when related items are enhanced or weakened. Another is to predict how these principles lead to larger scale memory phenomena. Here we discuss the limitations of existing models and propose SAMPL: the Spreading Activation and Memory PLasticity model, a unified account of the malleability of memory.

Retrieval enhances memory for practiced items, while unpracticed but related items can undergo either forgetting (e.g., retrieval-induced forgetting, RIF)^1–4^ or enhancement (e.g., the context repetition effect, statistical learning of predictive associations^2^). Typically, memory models either focus on studies of forgetting of related items or enhancement, but not both.

In the first group, one biologically plausible proposal for explaining forgetting phenomena, such as RIF, is a neural network model using oscillating inhibition^3^. However, this model has not been shown to explain the enhancement of related but unpracticed items, is computationally expensive, and has so far only been tested on small memory networks. Another proposed explanation of RIF is the non-monotonic plasticity hypothesis (NMPH)^10^. This approach applies a “rich get richer, poor get poorer” update rule that shapes activation dynamics, enhancing associations between strongly activated items and weakening associations between moderately activated items, whether or not they were practiced. However, the NMPH has not been applied to explain empirical findings on the enhancement of related items, nor has a computational model been proposed with the potential to explain NPMH effects at larger memory network levels, e.g., the network of memory relationships outside the items presented in a given experiment. It has been suggested that non-monotonic plasticity is the brain’s unsupervised learning rule for “housekeeping” of competition between memories at the larger network level.^4^ If this is the case, then a non-monotonic updating rule should in principle lead to a wide range of human memory behavior including both enhancement and forgetting effects. However, no model so far has incorporated both non-monotonic plasticity and the spread of activation across memory networks to test whether these two principles alone are sufficient to account for complex memory phenomena.

In the second group, models for learning predictive representation^5–7^ have been proposed to explain phenomena involving enhancement of related items, e.g., in context repetition and statistical learning studies. Whether as neural network models^3^, reinforcement learning and representation learning models^8^, or graph network models, these approaches take into account the larger network structure of associations. However, they have not been shown to explain forgetting of related but unpracticed items.

We propose SAMPL: the Spreading Activation and Memory PLasticity model, a unified account of how memory retrieval leads to both memory enhancement and forgetting. SAMPL bridges principles from the above two groups of models, applying a non-monotonic plasticity updating rule across a relational memory network structure. SAMPL represents items stored in memory as nodes in a graph, and associations between items as weighted edges connecting those nodes.

SAMPL captures the plasticity of memory using two simple rules: (1) Spreading activation. Retrieving an item initiates a cascade of activation that spreads across the graph to connected nodes, activating related items that were not directly remembered^1^. This gradually decaying cascade of activation can reach memories that are multiple associations away in the larger network. (2) Non-monotonic plasticity^3, 4^. Edge weights are updated, strengthening edges between strongly activated nodes and weakening edges between moderately activated nodes.

We validate SAMPL by comparing its output to human behavior in simulations of human memory experiments^2, 9, 10^. We simulate three representative memory phenomena: retrieval-induced forgetting, the context repetition effect, and memory alignment in multi-person conversational networks. These simulations test two hypotheses. First, we test the hypothesis that spreading activation and non-monotonic plasticity are sufficient for human-like memory enhancement and forgetting. Second, we test the hypothesis that non-monotonic plasticity (“rich get richer, poor get poorer”) is a better fit to human behavior than monotonic plasticity, where associations are adjusted proportionally to the activity level at each node.

The following simulation results show that the combination of spreading activation and non-monotonic plasticity implemented in SAMPL is sufficient to simulate both forgetting and enhancement of related items. In all simulations, non-monotonic plasticity produced human-like behavior while monotonic plasticity did not, showing that non-monotonic plasticity is important for explaining human memory phenomena.

## Methods

### The model

SAMPL consists of a memory network, a spreading activation rule, and an edge weight updating (plasticity) rule. The memory network is a graph where nodes represent memory items, edges represent connections between items, and edge weights represent the strength of those connections. When a set of nodes is activated, those nodes are assigned an activation value, and the spreading activation rule propagates that activation through the network, setting activation values for all connected nodes. Then the weight updating rule alters all edge weights based on the final activation values at each node. Detailed explanations of the spreading activation and weight updating rules follow, and a worked example is presented in Figure 3.

#### Spreading activation rule

Activation is propagated through the network using a recursively defined updating rule, such that the current activation state depends on the previous activations and edge weights. A simplified version of this updating rule is presented in equation (1). Let *a* be a column vector of activations, let *W* be the weight matrix of the memory network, and let γ be a discounting constant. Given an initial activation state *a_init_*, the final activation state *a_final_* after *t* propagation steps can be expressed as follows:

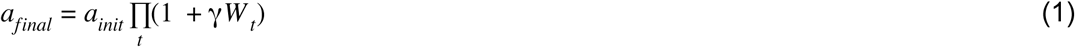

The complete form of the updating rule differs from the simplified version in equation (1) in three ways: Nodes only “fire” (i.e., spread activation to downstream nodes) when their activation is above the minimum threshold of the weight updating function, nodes that have fired are not allowed to fire again, and activations are restricted to the range [0, 1].

Here we provide a complete statement of the recursive updating rule for propagating activation through the network.

To only allow nodes to fire when their activation is above the minimum threshold of the NMP function, we define *f* (*a*) as the function that sets activations below that threshold to zero. Let *a* be a vector of activations with length equal to the number of nodes in the network *n*, indexed by *i* ∈ {1… *n*}, and let *x_min_*. be the smallest value in the domain of the NMP function that produces a non-zero output.

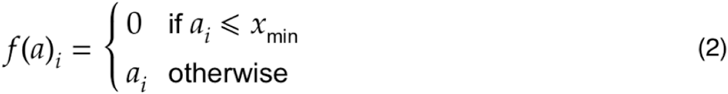

To prevent nodes that have fired from firing again, eliminating feedback loops in the network, we define *L*(*W*, *a*) as the function that sets a node’s inbound edge weights to zero if it will fire. Let *W* be the *n* × *n* weight matrix of the memory network, indexed by *i*, *j* ∈ {1… *n*}.

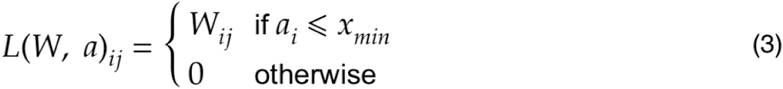

To restrict activation values to the range [0, 1], we define a clip function κ(*a*)

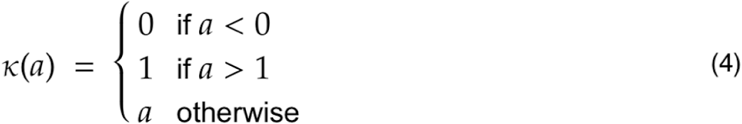

The complete statement of the spreading activation update rule has the same form as its simplified presentation in equation (1), but *f*, *L*, and κ are applied at each recursion step. Recursion terminates when all activations in *a* are 0.

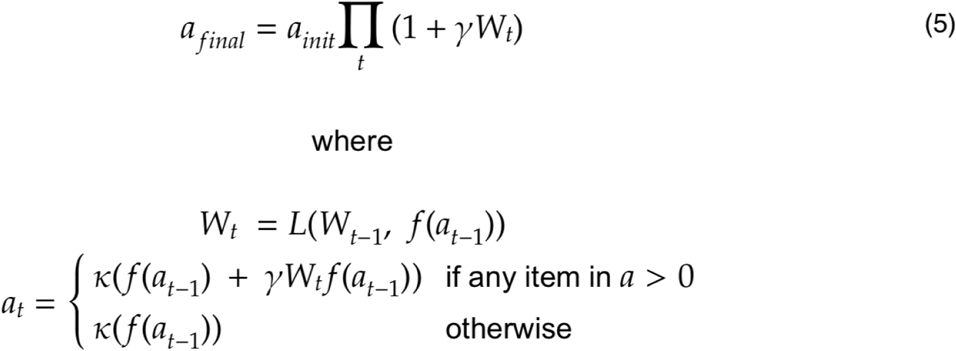

#### Weight updating rule

For each edge, the smaller of the two activation values at its corresponding nodes is used as input to a piecewise weight updating function, yielding an adjustment value which is added to the edge weight. The weight updating function has four parameters: suppression (y-min), enhancement (y-max), the width of the suppressive dip, and the center of the suppressive dip, visualized in Figure 2.

**Figure 1.**
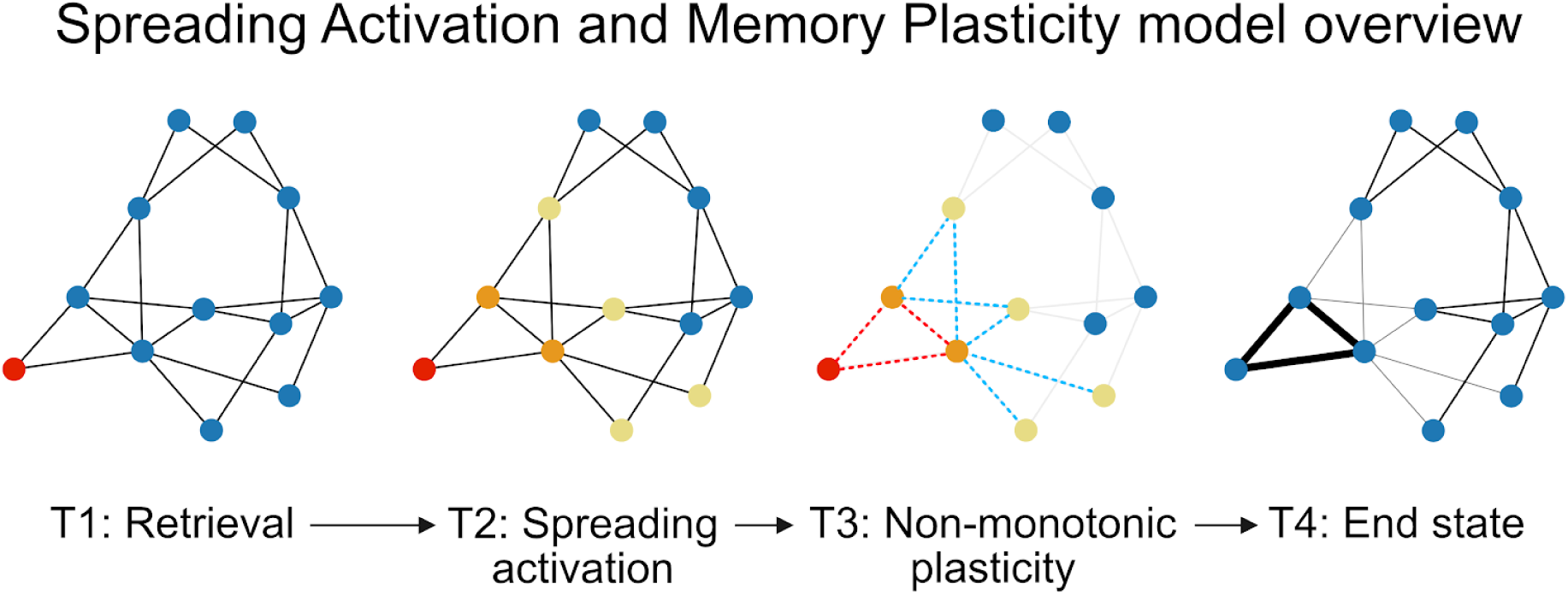
SAMPL: Spreading Activation and Memory PLasticity model schematic. (T1) Memory retrieval activates a node in the memory network (red node; colors denote the level of activation). (T2) This activation is spread across the graph, decaying according to the edge weights and the discount parameter (orange and yellow nodes). (T3) Edges are reweighted non-monotonically corresponding to the level of activation of their nodes: High activation strengthens connections (dashed red edges), moderate activation weakens connections (dashed blue edges), and low activation leaves connections unchanged (light grey edges). (T4) The final state of the memory network reflects changes in edge weights (compare to T1).

**Figure 2.**
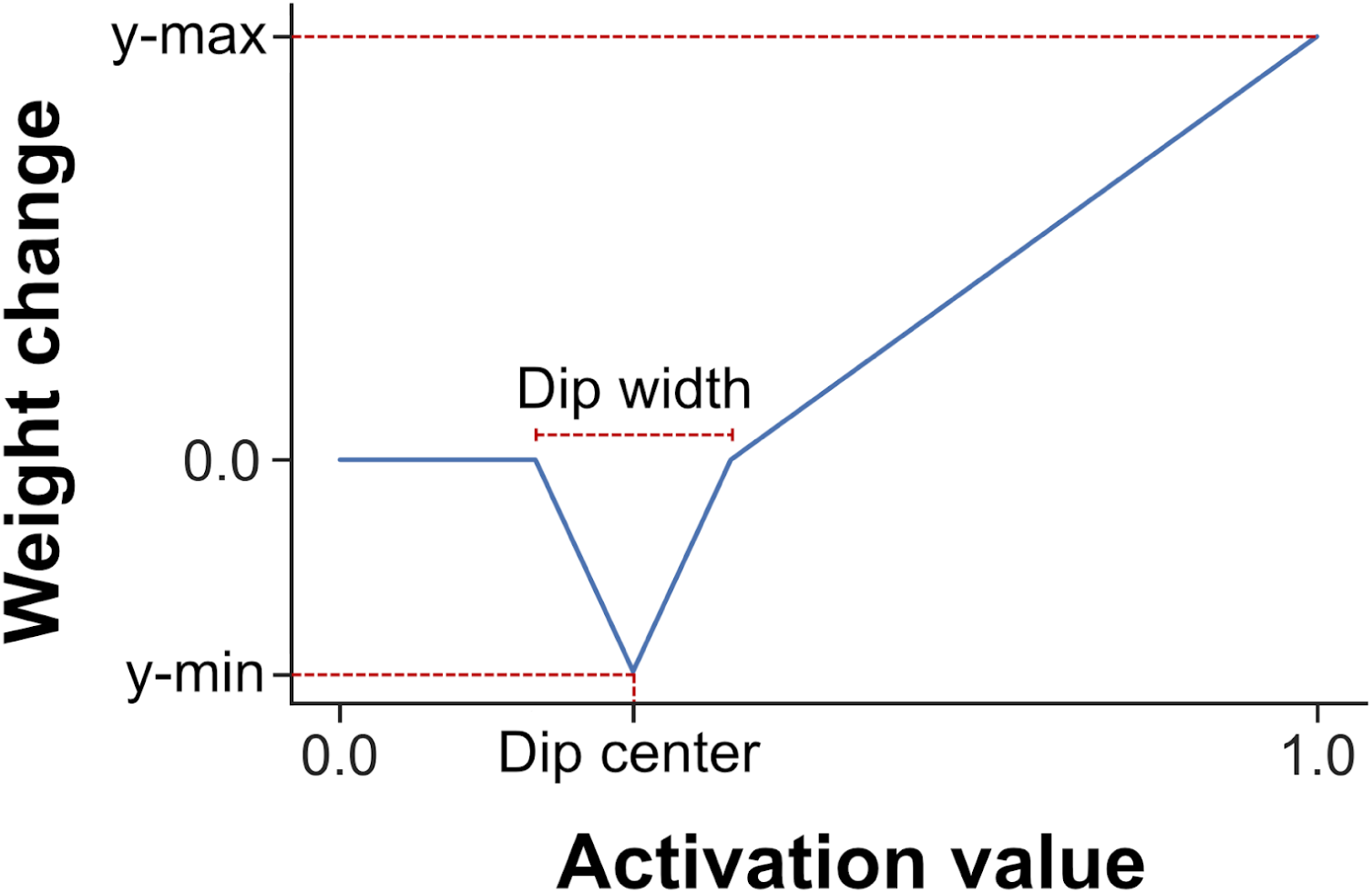
Non-monotonic weight updating function. The SAMPL edge weight updating function implements a non-monotonic plasticity (NMP) rule. Edge weights between two nodes increase when they are strongly activated, decrease when they are moderately activated, and do not change when they are weakly activated. The shape of the weight updating function is determined by four parameters: suppression (y-min), enhancement (y-max), the width of the suppressive dip, and the center of the suppressive dip (which determines the range of moderate activation, in which case edge weights are decreased). As such, the suppressive dip enables non-monotonic plasticity. Note that if the suppression (y-min) parameter is zero, then the function becomes monotonic.

SAMPL can accommodate a wide range of experimental paradigms. For example, activation values can be interpreted as the probability that an item will be remembered, or a threshold can be applied such that only nodes with higher activation will be remembered. Similarly, connections and edge weights can be set based on experiment-specific details. For example, an edge weight could indicate how often two items co-occur in a text corpus, or could be set to reflect behavioral relatedness norms. In Studies 1, 2, and 4, SAMPL was adapted to existing paradigms from the memory literature. In Studies 3 and 4, SAMPL was used in agent-based modeling simulations, where each agent uses an independent instance of the model as its internal memory. In Study 4, the model was extended to include separate episodic and long-term memory stores.

## Simulations

### Study 1: Retrieval-induced forgetting

#### Methods

To assess whether SAMPL produces retrieval-induced forgetting (RIF), we simulated experiment 1 from Anderson, Bjork, & Bjork (1994)^9^, henceforth ABB (1994). See Supplementary Figure 1 for a schematic of RIF experiments and Supplementary Table 1 for the items used in the ABB (1994) study. In this experiment, participants first studied associations between category labels and category exemplars. Participants then engaged in retrieval practice of half of the exemplars from a subset of the categories. During a final memory test, participants were asked to engage in free recall of exemplars from a single category for 30s. The study found that exemplars that were not practiced but belonged to a practiced category (RP- items, see below) were remembered less frequently than exemplars that were not practiced and belonged to an unpracticed category (NRP items). The authors called this effect retrieval-induced forgetting. Following the convention established by ABB (1994), here we use the shorthand *RP+* to indicate practiced items, *RP-* to indicate unpracticed items from a practiced category, and *NRP* to indicate unpracticed items from an unpracticed category.

We created a memory network with nodes corresponding to the words used in ABB (1994). To ensure realistic word-to-word associations, edge weights were determined using a separate semantic embedding model^11^ trained on the Google News corpus, a large collection of English language news reports. Specifically, the edge weights between nodes in the memory network were set to the cosine distances between the corresponding word vectors in the semantic embedding model. (N.B.: The “strong” high taxonomic frequency vs. “weak” low taxonomic frequency category distinction made in ABB is not well-captured by distances between word vectors in the semantic embedding model, so we did not attempt to replicate this aspect of their analysis.)

To simulate the study phase of experiment 1 from ABB (1994), edge weights from category labels to exemplars were set equal to the first value of the weight updating function above the suppressive dip. This ensured that simultaneous activation and weight updating would strengthen connections between category labels and exemplars. Categories were selected for retrieval practice in a counterbalanced order, as in ABB (1994). To simulate retrieval practice, each RP+ item (i.e., practiced exemplar) was activated at the same time as its corresponding label, activation was spread across the network, and edge weights were adjusted according to the weight updating function. To simulate final testing, the final item-label-to-category edge weight was used as a proxy for the proportion of successful recalls. Model parameters were set based on the results of a grid search (see Supplementary Materials for details), where the search cost of each parameter set corresponded to the sum of the differences between the model results and the results from ABB (1994).

### Study 1 Simulation Results

SAMPL produced a retrieval-induced forgetting effect (Figure 4). Both RP- and NRP items were remembered less frequently than RP+ items, and RP- items were remembered less frequently than NRP items [ABB (1994) difference: -.11, simulation difference: -.05, Welch’s t(142)=-15.39, p<.001] (Figure 4, top left). The best parameter set found in the grid search included moderate enhancement (y-max=.1), a small suppressive dip (y-min=-.01, dip center=.3, dip width=.35), and a moderate discount (γ=.3) (Figure 4, top right). Inspection of the edge weight matrix after simulation showed that the edge weights between category labels and RP+ items were maintained while the edge weights between category labels and RP- items were weakened, without any large changes involving NRP items or the rest of the network (Supplementary Figure 1). Parameter settings with more suppression or enhancement produced results less similar to ABB (1994), although a wide range of discount (γ) values appeared viable (Figure 4, middle). Notably, we also tested monotonic plasticity update rules (plasticity functions with no dip), comparing simulation outcomes. Using a monotonic weight updating function with no suppressive dip eliminated the retrieval-induced forgetting effect (difference: 0) (Figure 4, bottom).

**Figure 3.**
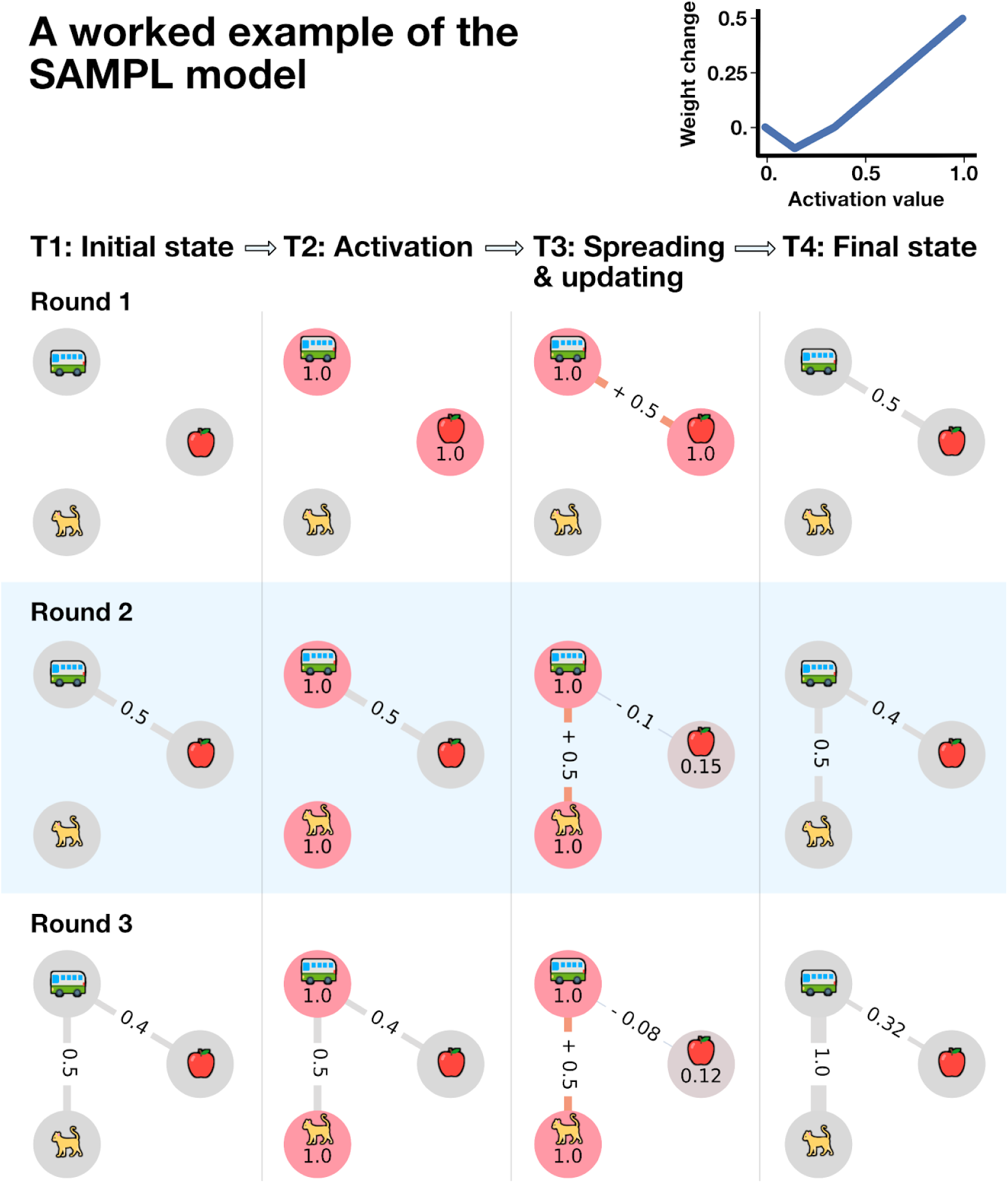
A worked example of the SAMPL model. In SAMPL, we model items in memory as nodes in a graph. When one or more items are retrieved, activation spreads across the graph—according to the strength of associations between items with cascading decay—and edge weights are updated. This example uses the non-monotonic weight updating function in the upper right corner of the figure, with discount factor γ = .3, in three rounds of retrieval. (Round 1) Initially, there are three items in memory (apple, bus, and cat). Let us assume the edge weights between them are all zero. The apple and bus nodes are activated simultaneously. Because all edge weights are zero, no activation is propagated to the cat node. The *smaller* activation value of both nodes on the apple-to-bus edge is 1.0, so the edge weight is adjusted by the non-monotonic plasticity rule to .5. (Round 2) The bus and cat nodes are activated simultaneously. The activation that spreads to the apple node is equal to the activation at the bus node (1.0) scaled by the edge weight between the two items (.5), as well as the discount factor (γ = .3), or 1 * .5 * .3 = .15 . Since the activation at both the bus and cat nodes is 1.0, the edge between them is incremented by the update rule to .5. By contrast, the minimum activation of the bus and apple nodes is 0.15 (at the apple node). This moderate activation level is in the suppressive dip of the weight updating function (top right). Therefore, the update rule adjusts the edge weight by -.1, leading to retrieval-induced forgetting. (Round 3) Simultaneously activating the bus and cat nodes a second time results in further reinforcement or enhancement of the connection between them. However, note that the weight updating rule leads to the further weakening of the edge weight from bus to apple. *Here we use an NMP function compared to* Figure 1 *where only 0 activation leads to no weight update*.

**Figure 4.**
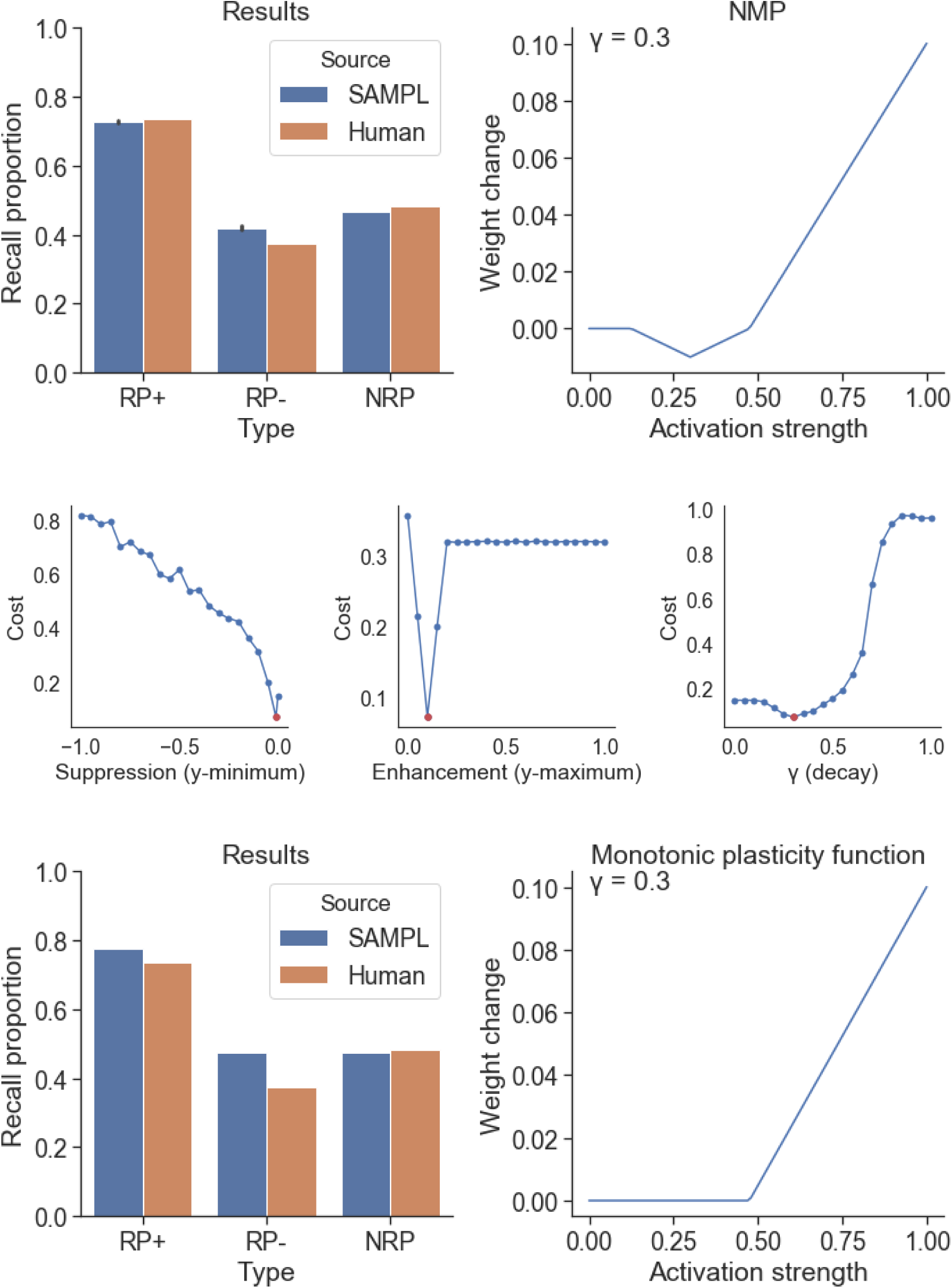
SAMPL simulates retrieval induced forgetting. (Top left) SAMPL simulation results compared to human behavior reported in Anderson, Bjork & Bjork (1994), or ABB (1994). SAMPL produces behavior to human results. (Top right) Optimal plasticity function and discount (γ) parameter values determined by grid search. (Middle) The fit between SAMPL and human results changes as a function of the suppression, enhancement, and discount (γ) parameters, as other model parameters are held constant. Lower cost values (Y axis) indicate a better fit of the behavior of SAMPL to ABB (1994) results. The red dot indicates optimal parameters found by grid search. (Bottom) When the suppressive dip is removed from the plasticity function (right), retrieval-induced forgetting does not occur, i.e., the blue bars in the RP- and RP+ condition are not different (left).

### Study 2: The context repetition effect

#### Methods

To assess whether SAMPL produces the context repetition effect, we conducted a simulation study replicating experiment 1 from Smith, Hasinski, & Sederberg (2013), henceforth SHS (2013). Participants in SHS (2013) viewed a series of images organized in a triplet structure, with the first two items in each triplet acting as the context for the third item, called the target. This study design supported measurement of both the *item repetition effect* and the *context repetition effect.* The *item repetition effect* refers to enhanced memory for targets that were repeated, while the *context repetition effect* refers to enhanced memory for targets whose context items were repeated, even if the target was not repeated. SHS (2013) distinguishes between a *predictive context effect,* where the context was repeated only *after* it was paired with the target (called the Target 1 context repetition effect in SHS 2013), and a *non-predictive context effect,* where the context was repeated *before* it was paired with the target (called the Target 2 context repetition effect in SHS 2013). Importantly, SHS (2013) found a moderate predictive context effect, but no non-predictive context effect. Retrieval of context items enhanced memory only for targets with which they were *previously* paired. This suggests the possibility that context enhancement is driven not simply by better memory for the context items, but by activation of the context items spreading across and reinforcing connections to target items they were paired with in the past.

We simulated the context repetition paradigm using SAMPL. Because the stimuli used by SHS (2013) were novel images assumed to have no pre-existing associations with one another, we set the initial edge weights of the graph to zero. To simulate the learning phase of the experiment, we generated a study list following the procedure described by SHS (2013). Each study list contained four types of triplets, organized in a two-by-two design: repeated context with repeated target; repeated context with novel target; non-repeated context with repeated target; non-repeated context with novel target. Study lists were constructed by randomly selecting 4 triplets of each type, for a total of 72 items and 96 item presentations (due to item repetition). Items were presented to the model by iterating over each study list using a 3-item rolling window. At each step, the three items in the window were activated, that activation was spread across the memory network, and edge weights were adjusted according to the weight updating function. Critically, the rolling window was allowed to extend across inter-triplet boundaries, mimicking the serial presentation approach of SHS (2013). To simulate the item-recognition test, the in-degree of each node was interpreted as a measure of relative memorability.

Model parameters were set based on the results of a grid search (see Supplementary Materials for details). The search cost function was designed to capture the important features of the result from SHS (2013): high item repetition effects, moderate context repetition effect for predictive context, and near zero context repetition effects for non-predictive contexts on a target item. Accordingly, the search cost of each parameter set was defined as the difference between the ratio of the item repetition effect to the predictive context repetition effect in the simulation and the same ratio in SHS (2013), plus one if a predictive context effect was *not* detected, and plus one if a non-predictive context effect *was* detected. The ratio was used because the simulation results used different units from SHS (2013).

#### Study 2 Simulation Results

The model produced item repetition effects and context repetition effects similar to SHS (2013) (Figure 5), including a strong item repetition effect, a moderate predictive context repetition effect, and no non-predictive context repetition effect. The predictive context repetition effect was smaller than the item repetition effect [Welch’s t(2158)=-26.57, p<.001] and greater than the non-predictive context effect [Welch’s t(1438)=29.84, p<.001] (Figure 5, left and middle). The best parameter set found in the grid search included both moderate enhancement (y-max=.4), a moderate suppressive dip (y-min=-.5, dip center=.2, dip width=.1), and a moderate discount (γ=.3) (Figure 5, right).

**Figure 5.**
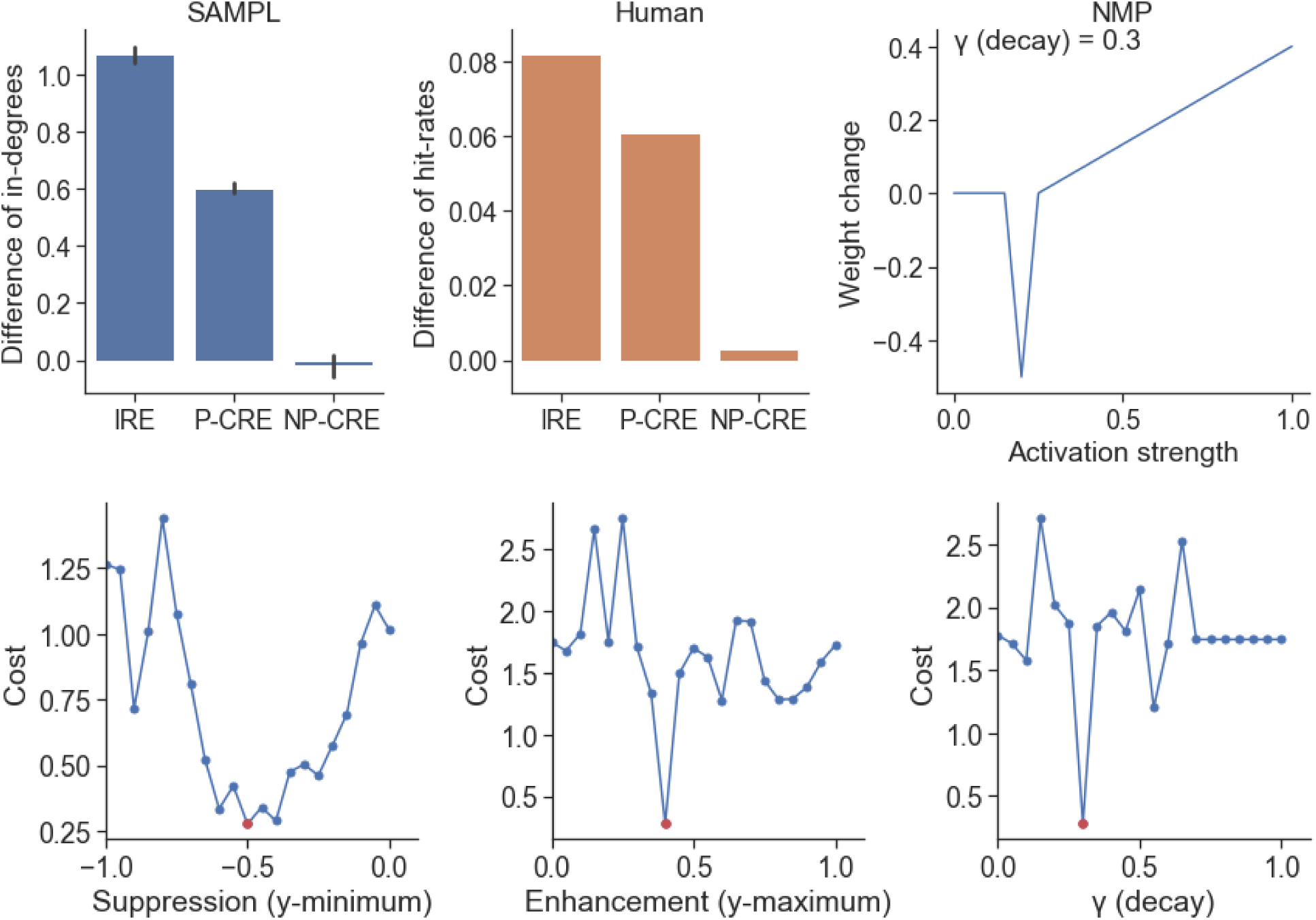
SAMPL simulates item repetition and context repetition effects. (Top) Simulation results (left) match human behavior (middle) reported in Smith, Hasinsky, Sederberg (2013), or SHS (2013). Repeating an item A, enhances the memorability of that item (IRE: item repetition effect). Repeating a predictive context for item A, i.e., a context that immediately preceded item A, enhances the memorability of the predicted item (P-CRE: predictive context repetition effect). Finally, repeating a non-predictive context for item A, i.e., a context that was previously presented immediately before a different item, B, does not enhance the memorability of item A (NP-CRE: non-predictive context repetition effect). Optimal plasticity function and discount (γ) parameters were determined by grid search (right). (Bottom) The fit between SAMPL and human behavior changes as a function of the suppression, enhancement, and discount (γ) parameters, as other model parameters are held constant. Lower cost values (y axis) indicate a better fit of the SAMPLE behavior to SHS (2013) results. The red dots indicate optimal parameters found by grid search.

We compared results using plasticity functions with different parameters. We observed that a broad range of suppressive dip values produced low search costs. However, given the best suppressive dip parameter value, even small changes to the enhancement and discount (γ) parameters resulted in rapidly increasing cost (Figure 5). Notably, using a monotonic weight updating function with no suppressive dip reduced the predictive context repetition effect and, interestingly, induced a non-predictive context repetition effect not observed in SHS (2013). Both the predictive and non-predictive context repetition effects were inversely correlated with the magnitude of the suppressive dip [Predictive: Pearson’s R=-.32, p<.001; non-predictive: Pearson’s R=-.27, p<.001], with the optimal suppression parameter (y-min=-.5) resulting in a non-predictive context repetition effect close to zero.

Importantly, this shows that non-monotonic plasticity plays a counterintuitive but critical role in context enhancement. NMP prevents generalization of an item to non-predictive contexts because it limits associations between novel items and previously presented contexts.

### Study 3: Memory synchronization in communicating agents

#### Methods

Memory enhancement and forgetting effects including retrieval-induced forgetting and the context repetition effect have been observed in studies of dyadic as well as multi-agent conversation^10, 12, 13^. We assessed whether SAMPL can simulate the synchronization of remembered and forgotten items in conversational networks^10, 12^. Before testing the model in a network of multi-agent simulations, we conducted a dyadic agent-based modeling study. This study measured the effect of different conversation strategies on memory similarity across two agents. Because the focus of this study was to test the effect of different conversation strategies, a single non-monotonic weight updating function with moderate enhancement (y-max=.15), a small suppressive dip (y-min=.025, dip center=.3, dip width=.4), and a moderate discount (γ=.3) was used. Both agents were given the same vocabulary by assigning the same labels to their graph nodes. This vocabulary contained a range of strongly and weakly associated items. As in Study 1, to ensure realistic word-to-word associations, edge weights were determined by taking the cosine distance between words in a word2vec model^11^ trained on the Google News corpus.

We operationalized conversations as series of exchanges. In each exchange one agent, designated *the sender*, selected a pair of words to be activated in the other agent, designated *the receiver.* The word selection process is described below. This activation was then spread across the receiver’s memory network, and the receiver’s edge weights were adjusted according to the weight updating function. The process was then repeated with the sender and receiver roles reversed, so each agent acted as both sender and receiver in each exchange.

Mnemonic convergence (or synchronization) increases when the agents’ memory network graphs become more similar to one another, as measured by the correlation distance between the edge weight matrices of the two agents after every exchange. We simulated 100 conversations, each containing 50 exchanges, supporting the calculation of a mean distance over time trajectory. For one agent, the edge weights were shuffled at the start of each conversation, ensuring that the agents had different but comparable memory networks.

We compared three conversation strategies used by the sender to select a word pair. For the *uniform* strategy, both words were selected from the sender’s vocabulary uniformly at random without replacement. For the *mean in-degree* strategy, the mean in-degrees of all of the words in the sender’s vocabulary were normalized to create a probability distribution, and both words were selected by taking random draws without replacement from that distribution. In the *paired associate* strategy, the first word was selected with the mean in-degree strategy, and the second word was selected from the associates of the first word. That is, the second word was selected by normalizing the outbound edge weights from the first word to create a probability distribution and taking a random draw from that distribution. The paired associate strategy was designed to resemble human conversations, where related words tend to co-occur.

#### Study 3 Simulation Results

We found that changes in mnemonic convergence strongly depended on conversation strategy (Figure 7). For all strategies, the initial correlation distance between the agents’ edge weight matrices was approximately equal [Uniform: initial distance=.98, 95% CI [.93, 1]; Mean in-degree: initial distance=.99, 95% CI[.96, 1.02]; Paired associate: initial distance=.97, 95% CI [.93, 1]]. Each strategy produced different convergence behavior. Uniform selection resulted in no change in correlation distance [Δ distance=0, final distance=.98, 95% CI[.97, 1.03]]. Mean in-degree selection resulted in some reduction in distance [Δ distance=-.09, final distance=.9, 95% CI [.87, .93]] that was greater than the reduction for uniform selection [Welch’s t(198)=-.3.44, p<.001]. Paired associate selection resulted in a large increase in similarity, or reduction in distance [Δ distance=-.79, final distance=.19, 95% CI[.18, .21]] that was greater than both the reduction for uniform selection [Welch’s t(198)=-36.68, p<.001] and for mean in-degree selection [Welch’s t(198)=-31.72, p<.001].

**Figure 6.**
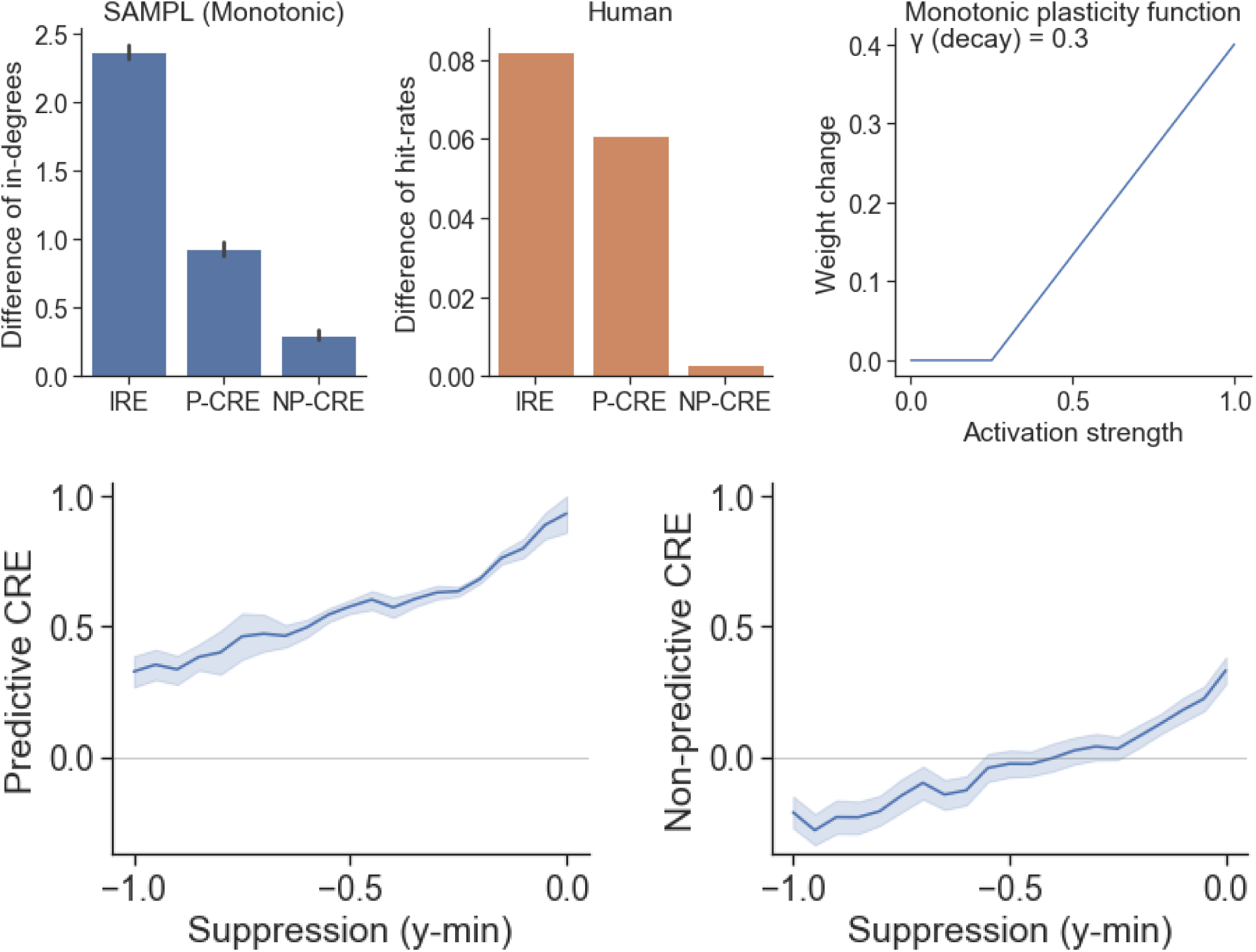
The role of non-monotonic plasticity in context repetition effects. (Top) Removing the suppressive dip from the plasticity function, i.e., using a monotonic weight updating rule (right) reduced the predictive context repetition effect (left) and, interestingly, induced a non-predictive context repetition effect not observed in human behavior in SHS (2013). (Bottom) Both the predictive and non-predictive context repetition effects were inversely correlated with the magnitude of the suppressive dip. Importantly, this shows that suppression prevents generalization to non-predictive contexts.

**Figure 7.**
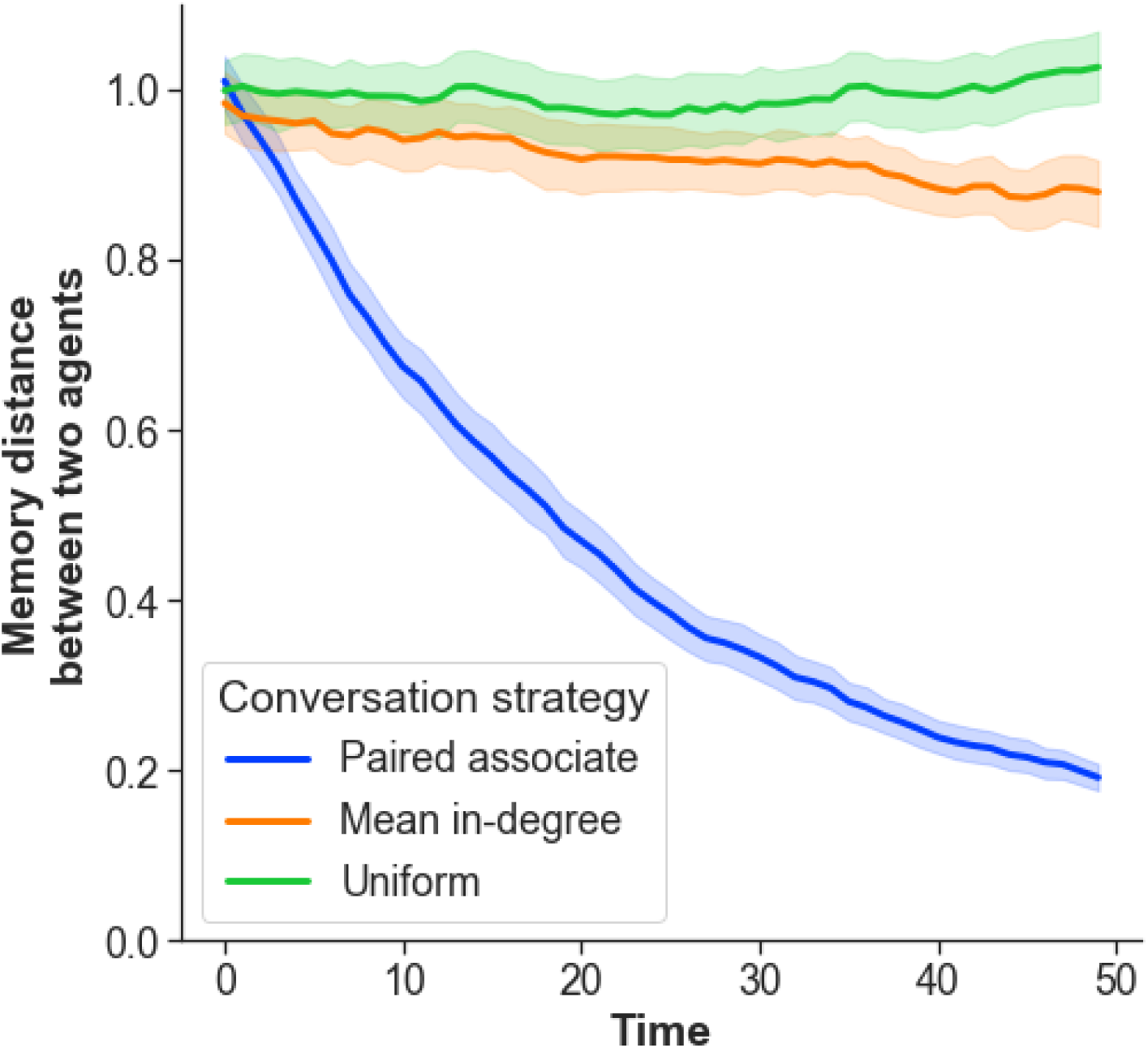
Memory networks become more similar during conversations. We used SAMPL to simulate conversations between two agents with different memory networks drawn from the Google News corpus. We tested how three different conversation strategies affected changes in their memory similarity as a function of their conversations over time, measured using correlation distance. *Uniform strategy*: For every round of conversation, the sender selected two words randomly, with uniform probability. The nodes corresponding to these words were activated in the receiver’s memory graph, which was then updated by SAMPL. *Mean in-degree strategy*: Same as the uniform strategy, except instead of selecting words from a uniform distribution, the probability distribution for selecting words was weighted by the mean in-degree of each word. *Paired associate strategy*: In this strategy the first word was selected using the mean in-degree strategy, and the second word was an associate of the first word, selected with a probability proportional to the strength of the edge weight between the two words. Here correlation distance over time is displayed on the Y axis for the three conversation strategies over the course of 50 exchanges. Non-monotonic plasticity created mnemonic convergence when agents stay “on topic,” i.e., when they exchanged words using the paired associate strategy, but not when they had random conversations.

Notably, in agents with the paired associate selection strategy the initial standard deviation of the edge weights was similar to the final standard deviation [initial mean std=.15; final mean std=.18]. This indicates that the observed mnemonic convergence was not simply convergence to a single value, and was not the result of floor or ceiling effects. Typical initial and final edge weight matrices are visualized in Supplementary Figures 2 and 3. These results indicate that SAMPL is sufficient to simulate the convergence of memory in communicating agents, but only when conversation stays “on topic,” consisting of exchanges of associated words.

### Study 4: Social networks and collective memory

#### Methods

To assess whether SAMPL is sufficient to simulate collective memory effects at the level of multi-agent networks, we conducted an agent-based modeling study replicating Coman, Momennejad, Drach, & Geana (2016)^10^, henceforth CMDG (2016). In the study phase of CMDG (2016), participants studied four facts about four fictional characters. In the pre-conversational recall test phase, participants were given a character name as a cue, and asked to recall the studied information. In the conversational recall phase, participants were split into 10-member communities. Each participant then had a series of dyadic conversations with partners from the same community, during which they were instructed to jointly recall the studied information. Each community was either assigned to a clustered condition or a non-clustered condition, determining the order in which participants conversed.

In the clustered condition, the network structure of the community contained two subclusters with a moderate global clustering coefficient (C=0.4), whereas in the non-clustered condition, the network structure consisted of a single large cluster (C=0) (Figure 8). Finally, in the post-conversational recall phase, participants were cued with character names and asked to recall facts about each character. CMDG (2016) assessed the mnemonic alignment between every pair of participants in each community, expressed as the change in memory similarity from pre- to post-conversation. They found that mnemonic alignment was higher for pairs of participants that were closer to each other in the social network. CMDG (2016) also assessed mnemonic convergence across entire networks, expressed as the average of pairwise memory similarity scores within each community. The results showed slightly increased convergence in the non-clustered network both pre- and post-conversation.

**Figure 8.**
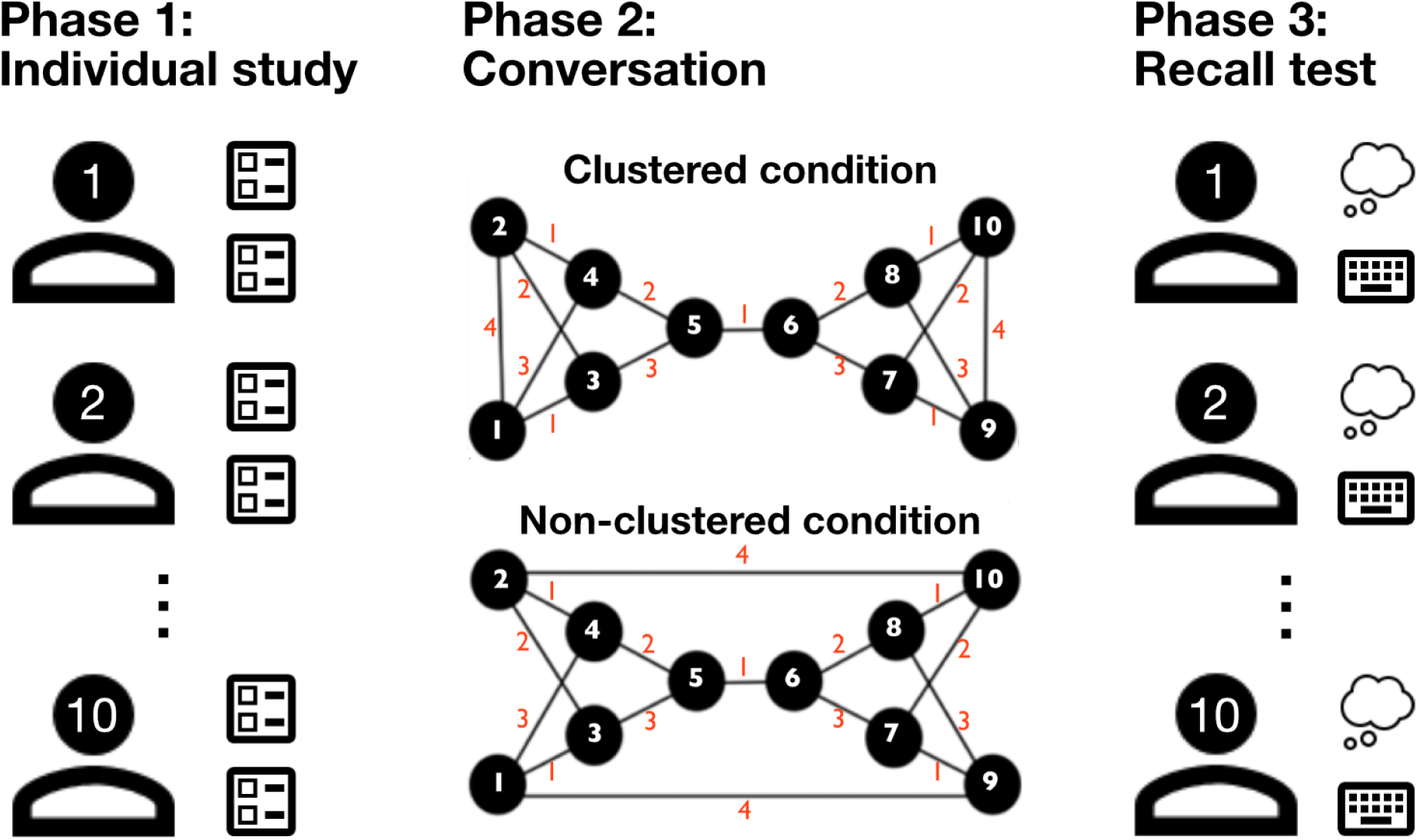
Design schematic for collective memory study. In Coman, Momennejad, Drach, Geana (2016), or CMDG (2016), participants first studied memory material and took a memory recall test (Phase 1). Then they each had 3 sequential conversations with others, determined by one of two social network graphs, clustered or non-clustered, each with 10 participants (Phase 2). Finally, they took a memory recall test (Phase 3). Comparing memory similarity, i.e., correlation, before and after the conversations revealed the effect of conversation on mnemonic convergence. In our simulation of this study, SAMPL agents were initialized with a memory network containing 20 nodes, corresponding to four characters and four facts about those characters, as in CMDG (2016). (Left) In phase 1, individual study was simulated by randomly selecting and sequentially activating half of the node pairs in each agent’s network, and applying SMPL. This ensured that each agent had memory for some, but not all facts about the characters. (Center) In phase 2, pairs of agents (represented by black circles) conversed in an order dictated by whether they were in the clustered or non-clustered network condition (order shown in red numbers). Conversational exchange proceeded using the paired-associate strategy and SAMPL. (Right) In phase 3, the recall test was simulated by averaging pairwise correlations between the final memory networks of the agents.

We simulated the collective memory paradigm of CMDG (2016) using an agent-based modeling approach similar to that of Study 3. Agents were initialized with a memory network containing 20 nodes, corresponding to four characters and four facts per character. Because participants in CMDG (2016) began with no knowledge of the fictional characters, all edge weights in the agents’ memory graphs were initially set to zero.

The study and pre-conversational recall phases were simulated by a single study phase, designed to provide each agent with memory for some, but not all, facts about the characters, and to ensure that each agent remembered a different set of facts. For each agent, half of the character-fact node pairs were randomly selected, then each pair was activated in series, causing non-monotonic reweighting of the agent’s memory graph.

The conversational recall phase was simulated first by selecting pairs of agents in the order described in Figure 8, then by simulating a conversation between those agents using the paired-associate strategy as described in Study 3 above. Each conversation consisted of 100 paired-associate exchanges, including non-monotonic adjustment of each agent’s edge weight matrix. The agent architecture was elaborated to account for the episodic nature of the conversation paradigm. At the beginning of each conversation, the agent’s memory graph was copied, and changes during that conversational episode were applied only to the copy. At the end of each conversation, the original memory graph was replaced by a linear interpolation between the original graph and the copy, with a learning rate parameter controlling the amount of interpolation (from 0% to 100%).

To simulate the post-conversational recall phase, the correlation between the edge weight matrices of a pair of agents was used to measure memory similarity. This memory similarity measure was used to measure mnemonic alignment and mnemonic converge analogously to CMDG (2016). However, because we directly measured agents’ edge weight matrices rather than testing remembered and forgotten items, our results use different units and are on a different scale.

We used a simple grid search to find model parameters that best reproduced the main findings of CMDG (2016). The cost of each simulation at each point on the parameter grid was a weighted average of four values: the absolute error of the proportion-adjusted mnemonic convergence results, the absolute error of the proportion-adjusted mnemonic alignment results, the absolute error of the effect of network hops on alignment, and one minus the signed alignment value at one network hop. The first two values were proportion adjusted because our operationalization of convergence and alignment produced results on a different scale than the behavioral results, as described above. The fourth value, one minus signed alignment, was included for the same reason. The weights for these values were 1, 3, 2, and 3, respectively, prioritizing results that matched the alignment finding of CMDG (2016) even if they differed slightly in absolute terms.

#### Study 4 Simulation Results

The model reproduced the main findings of CMDG (2016): Mnemonic convergence was higher post-conversation than pre-conversation [Clustered: t(158)=59.85, p<.001; Non-clustered: t(158)=82.87, p<.001], and post-conversation convergence was slightly higher in the non-clustered condition [t(78)=3.12, p=.002] (Figure 9, top). Simple linear regression confirmed that mnemonic alignment decreased with network distance both in the clustered condition [F(1, 3598)=1198, adj. R^2^=.25, β_distance_ =-.11, p<.001] and the non-clustered condition [F(1, 3598)=496.8, adj. R^2^=.12, β_distance_ =-.14, p<.001] (Figure 9, bottom). The best parameter set found in the grid search included moderate enhancement (y-max=.5), a relatively large suppressive dip (y-min=-.8, dip center=.3, dip width=.4), a moderate discount (γ=.5), and a maximal learning rate (learning rate=1.0) (Figure 9).

**Figure 9.**
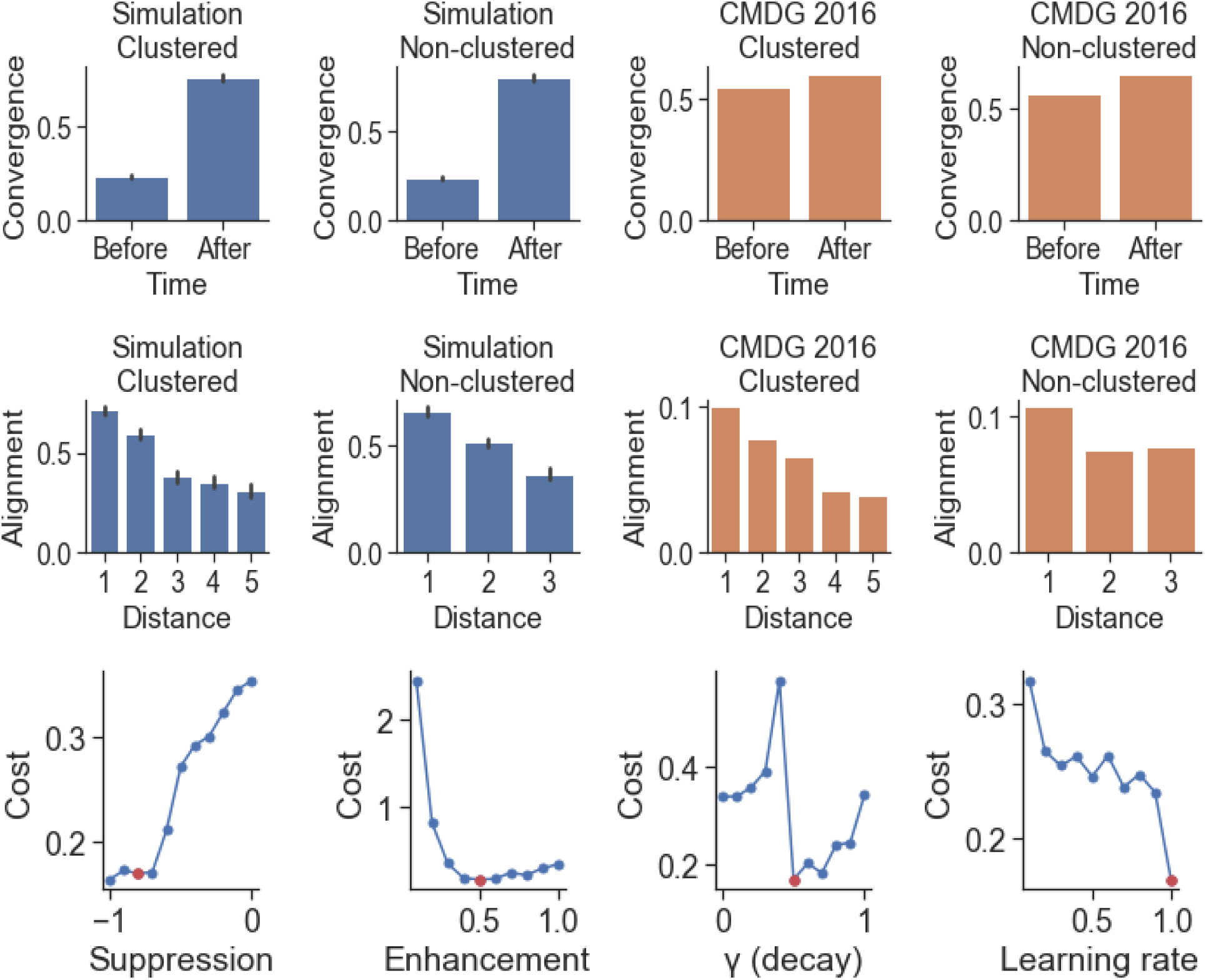
Agent-based modeling with SAMPL simulates memory convergence in social networks. (Top and middle) SAMPL simulation results (blue) matched human behavior from CMDG (2016) (orange). Mnemonic convergence was higher post-conversation, and post-conversational convergence was slightly higher in the non-clustered condition. Mnemonic alignment between two agents decreased with their degree of separation in the social network graph. (Bottom) The fit between SAMPL and human results changes as a function of the suppression, enhancement, discount (γ), and learning rate parameters, as other model parameters are held constant. Lower cost values (Y axis) indicates a better fit of SAMPL to CMDG (2016) results. Red dots indicate optimal parameters found by grid search.

Given relative strong suppression, a range of enhancement values produced low search cost, but small changes to the discount (γ) parameter resulted in increased cost (Figure 9). This suggests that small changes in how activation propagates through individual memory graphs has an effect on memory synchronization at the social network level. In particular, when the discount (γ) or suppression parameters were decreased, causing activation to propagate greater distances within individual memory graphs, then the simulation produced results diverging from the human results of CMDG (2016).

Importantly, using a monotonic weight updating function with no suppressive dip nullified the difference in convergence between the clustered and non-clustered conditions [t(158)=1.23, p=.22], as well as the effect of social network distance on alignment in both the clustered condition [F(1, 3598)=2.39, adj. R^2^=0, β_distance_ =0, p=.12] and the non-clustered condition [F(1, 3598)=3.77, adj. R^2^=0, β_distance_ =0, p=.05] (Figure 10). This suggests non-monotonic plasticity plays an important role in the formation of collective memory.

**Figure 10.**
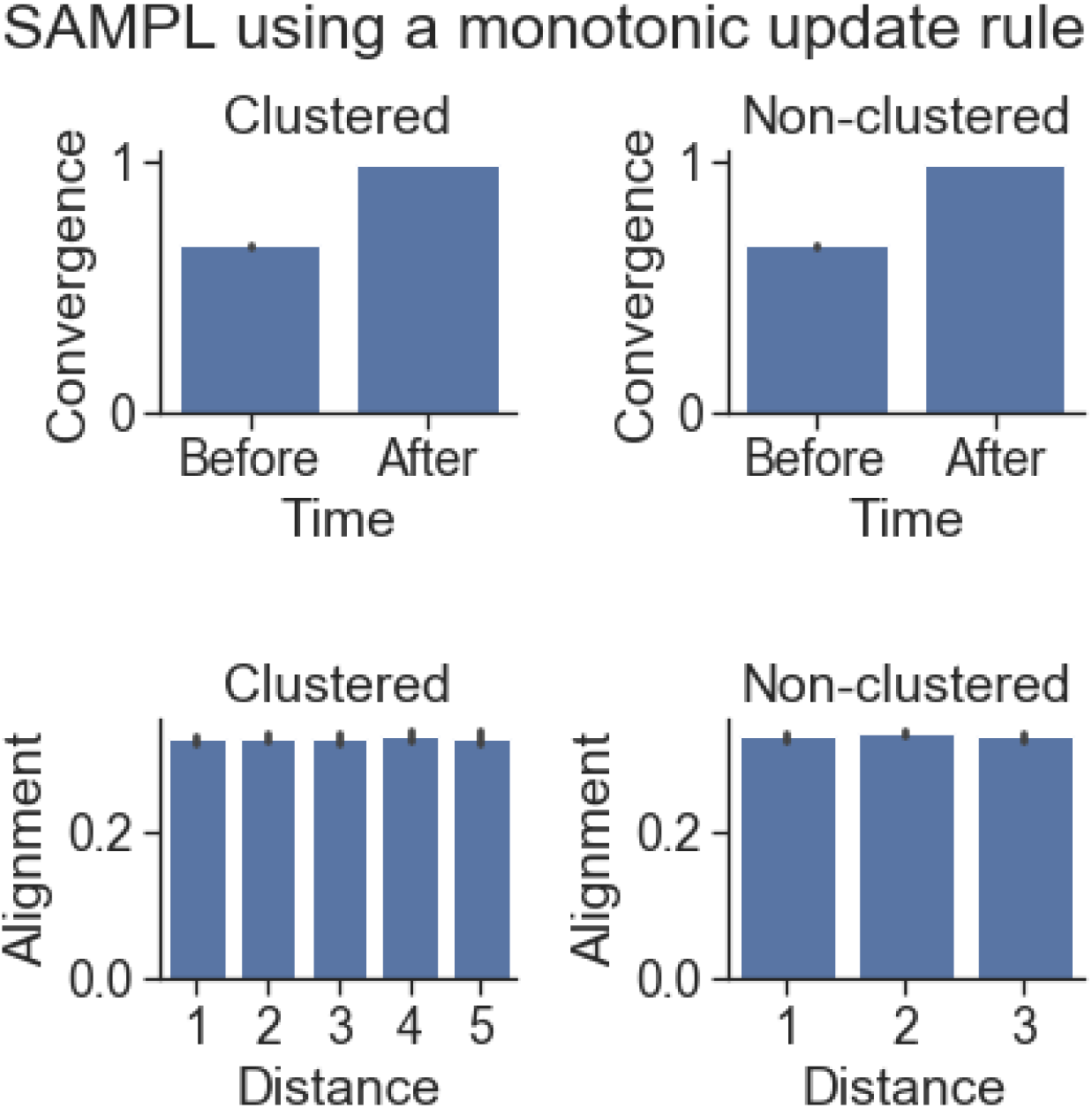
Monotonic plasticity cannot simulate empirical findings on mnemonic convergence. Using a monotonic weight updating function with no suppressive dip, instead of NMP, nullified both the observed difference in memory convergence between clustered and non-clustered conversational networks, as well as the overall effect of degree of separation between two agents on memory alignment. This shows that SAMPL requires a non-monotonic plasticity rule to simulate CMDG (2016).

## Discussion

We propose SAMPL, the Spreading Activation and Memory PLasticity Model. SAMPL is a graph-based model of how memory retrieval changes what will be remembered in the future. SAMPL combines two simple rules: a propagation rule for spreading activation in memory networks, and a plasticity rule for updating associations between memories. We hypothesized that spreading activation and non-monotonic plasticity would be sufficient to account for seemingly contradictory memory enhancement and forgetting phenomena. Confirming this hypothesis, we show that SAMPL simulates context-based memory enhancement^8^ and retrieval-induced forgetting ^9, 14^. Further, simultaneous support for enhancement and forgetting allows SAMPL to simulate the synchronization of memory in conversational dyads and in multi-person conversational networks.^9^

Our second hypothesis was that the memory update rule must have a non-monotonic shape. That is, while low activation would not lead to change, moderate activation would lead to weakening of associations between memories, and high activation would lead to strengthening of associations. To test this, we used a grid search to find the best fitting parameters for the update rule. This allowed us to compare the performance of update rules with and without a suppressive dip for retrieval-induced forgetting (Figure 4), the context repetition effect (Figure 6), and memory convergence in social networks (Figure 10). Confirming our hypothesis, we found that a non-monotonic “dip” in the plasticity curve was required to reproduce human results. Conversely, plasticity rules with no dip led to results that did not fit human behavior.

Importantly, SAMPL reveals that non-monotonic plasticity plays a counterintuitive but crucial role in context-based enhancement. We found that non-monotonic plasticity prevented generalization of an item to a non-predictive context. This is because NMP suppresses the association between novel items and previously presented contexts, in line with human behavioral findings in non-predictive context repetition studies^2^ (Figure 6). This finding has implications for understanding the role of memory processes in adaptive and maladaptive generalization, in both health and disease. It also informs the design of novel experimental paradigms to study generalization and test SAMPL predictions. In the long-run, we expect that SAMPL will help simulate computational interventions designed to correct maladaptive generalization.

### Relationship to existing memory models

In the past two decades, two groups of different but related models have been proposed to capture memory enhancement and forgetting phenomena. All of these models capture the enhancement of directly retrieved, or practiced, items. However, retrieval also induces changes in the memory of related items that are, importantly, unpracticed. While one group of models focuses on capturing the forgetting of these related items, another group focuses on the enhancement of related or predictive items. Our simulations capture both of these behaviors using simple propagation and plasticity rules.

The first group of models focuses on explaining forgetting of related unpracticed items. Non-monotonic plasticity alone predicts enhancement and weakening of associations depending on activation levels, but does not in itself specify how those activations propagate and change across memory networks. This requires specifying what items are in memory and the strengths of associations between them. SAMPL, the Spreading Activation and Memory PLasticity Model, addresses this limitation by combining a non-monotonic update rule and a spreading activation rule, both of which are applied to realistic memory networks, such as those derived from a large corpus of human data. Furthermore, previous studies have suggested that the NMPH explains retrieval-induced forgetting, but have not directly investigated enhancement of related but unpracticed items. Our results show that spreading activation and non-monotonic plasticity as implemented in SAMPL are sufficient to produce both forgetting and enhancement of related unpracticed items, providing a unified account of memory change.

Fitting the parameters of SAMPL to observed human behavior allows us to both estimate the optimal shape of the plasticity function and to track changes in each item’s activation over time. As such, SAMPL predicts both enhancement and forgetting across a wide range of tasks.

Another model for explaining retrieval-induced forgetting is a neural network model that implements an oscillating inhibition mechanism^3^. An advantage of this model is that it is mechanistic and biologically plausible. However, compared to SAMPL, its implementation is computationally expensive. In comparison, SAMPL operates on a higher level of abstraction, it does not require manual fitting of experimental items in the model, remains agnostic about how items are represented, and does not require a separate context element to explain context effects. Therefore, it can be easily adapted to a wide range of tasks and can predict behavior in larger networks at lower computational cost.

A second group of memory models focuses on memory enhancement due to associative and predictive relations among items. This includes models of association learning such as the successor representation^5, 8^, the Temporal Context Model^15^, the Context Maintenance and Retrieval model^16^, and complementary learning systems for statistical learning^17^. A series of recent empirical studies suggest the successor representation as a unifying principle for neural representations in memory organization and generalization at multiple scales^6, 7, 18, 19^. The memory enhancement study of context repetition effects simulated in the present paper suggests the successor representation as a potential account of their findings^2^. Furthermore, it has been shown that the temporal context model (TCM)^15^ is a special case of the successor representation, offering simple update principles for organization of and generalization across memory networks^5^. The Context Maintenance and Retrieval model (CMR) has been proposed as a generalization of TCM, separately accounting for semantic clustering (due to long-standing semantic associations) and temporal associations (due to study episodes)^16^. Note that both of CMR’s contributions are accounted for in SAMPL: semantic clustering is captured by the graph structure of the memory network, and temporal associations during an experiment are learned via sequential activation of study and practice items. Because these models are mathematically related^5, 15, 16^, here we discuss them in relation to one another.

Similarities among these models suggest that a unifying principle of memory representation and updating could parsimoniously explain the phenomena they capture. One proposal is that the successor representation might offer a unifying computational principle.^5^ However, while some models combine the successor representation with principles such as prioritized offline replay (SR-Dyna^6, 18^), the successor representation does not by itself constrain how sequential activation, memory search, and updating occur^19^. What is needed is a model that simultaneously offers principles for representation learning, memory search and retrieval, and memory update rules. Future modeling work is required for a direct mathematical comparison of these models.

SAMPL’s propagation is similar to the learning of the successor representation: both capture multi-step dependencies on a graph of association states, and this can be done by multiplying a matrix of association strengths by itself and scaled by a discount factor. SAMPL and SR differ in two ways. First, SAMPL implements a non-monotonic update rule while SR updates based on increases in predictivity. Second, SAMPL constrains multi-step dependencies by preventing loops in the spread of activation.

SAMPL is related to context-based models such as TCM and CMR in two ways. First, SAMPL captures temporal context: items that are activated and updated together serve as contexts for each other during future episodes. Because non-monotonic edge reweighting carries these updates forward in time, we anticipate SAMPL could simulate studies of temporal contiguity^20, 21^ effects^22, 23^ that are explained by TCM using sequential updating, as well as the successor representation^5^. Second, SAMPL captures semantic context: long-standing consolidated semantic associations are currently captured in edge weights that are determined using an independent corpus of data. This is comparable to the semantic clustering captured by CMR. SAMPL differs from context-based models in making the simplifying assumption that everything is an item: context is not modeled as a special and separate entity. Instead, context relationships are captured within SAMPL’s network relations and the temporal succession of a task’s episodes.

#### Relationship to brain function

We have shown that SAMPL, the Spreading Activation and Memory PLasticity Model, captures a range of human memory behaviors, but we have not yet shown how it may be implemented in the brain. To do so, we will first summarize related neural findings and how SAMPL corresponds to them. Neuroimaging and electrophysiological studies indicate that the hippocampus and prefrontal cortex (PFC) are the main components of the learning and memory network^24^, and that long-term and semantic memory networks are represented across the neocortex^25, 26^. Broadly speaking, a neural SAMPL would correspond to hippocampal and prefrontal processes.

Furthermore, it has been shown that the representations in the posterior hippocampus (analogous to dorsal hippocampus in rodents^27^) are at finer spatial, temporal, and predictive scales, while the anterior hippocampus represents dependencies at larger scales, allowing it to play a role in inference and abstraction^19, 28–30^. Compared to the anterior hippocampus, the prefrontal cortex may support even larger scales of abstraction, representing longer predictive horizons, enabling prospective memory, and inference over more distant associations^31–35^. Though the present version of SAMPL does not represent hierarchically organized networks at different scales of abstraction, applying different SAMPL parameters could simulate neural representations at different scales.

Assuming the brain implements the model, then SAMPL’s memory network component may be learned via hippocampal-prefrontal interactions and consolidated and stored in long-term memory throughout the neocortex^36^. Furthermore, since different brain regions support predictive representations at different scales, it is possible that the spread of activation discussed in the model reaches more distant memories in some brain regions compared to others. In the present model, spreading activation to more distant items in the memory network requires a lower discount factor. An important future direction is extending the model to capture the hierarchical organization of memory representations in the brain.

Neuroimaging studies often use representational similarity between two items as a proxy for the strength of association between those items^37–42^. For instance, say item A is retrieved and the subsequent spread of activation reaches a distal item G. A region with a large associative scale may show a high activation of G representations, in which case a non-monotonic plasticity update rule would increase the similarity between the representations of A and G. This is known as integration^4, 14, 40, 43–46^, generalization, or overlap^47^. Generalization can occur over short time-scales (during the experiment) and long time scales (after overnight consolidation). On the other hand, a region with a smaller associative scale may only show moderate activation of G, in which case non-monotonic plasticity would lead to a decrease in A-G similarity. This is known as separation^4, 46^. In regions with small enough scales where there is no G activation at all, the A-G connection remains unchanged.

By fitting the discount factor and plasticity function to data collected from different brain regions, it may be possible to estimate the associative scale of each region within and across brains. This may also allow us to explain seemingly contradictory findings showing that retrieval practice sometimes leads to integration and sometimes separation in the same brain region, for instance the anterior hippocampus. Let us assume that anterior hippocampus is a region in which activation can spread from X to Y. Given non-monotonic plasticity, X-Y integration takes place when Y is highly activated and separation takes place when Y is moderately activated. Therefore, the differences in integration and separation in the same region across studies might be due to different network relationships between X and Y: either different distances, strength of connections between X and Y, or a combination thereof.

In short, the associative scale of a given region determines whether and to what extent the spread of activation from retrieving one item activates another item. Non-monotonic plasticity can then explain why the same brain region can sometimes show integration and sometimes separation. Future research is required to identify how the associative scale of different brain regions relates to differences in parameters governing spread of activation and non-monotonic plasticity.

Here we have shown that non-monotonic plasticity always produced more human-like behavior than monotonic plasticity. However, both the optimal discount parameter and the optimal shape of the non-monotonic plasticity function differed for each study. This may be because different tasks demand different scales of memory organization. Earlier we discussed how different brain regions operate at different scales of the memory hierarchy. Furthermore, it has been suggested that the strength of inhibition and sparseness of representation in a brain region may determine the shape of NMP in that region^4^. For instance, hippocampal subregions with sparser activity and higher inhibition (CA3/DG) show separation and differentiation, and are hypothesized to have a different NMP shape from subregions with less sparse activity that show integration (CA1)^4, 17^. Furthermore, different discount factors (for a decaying spread of activation) and NMP shapes may be optimal for different real-world tasks, contexts, and goals. A previous study using functional magnetic resonance imaging (fMRI) has shown that the suppressive dip of the weight updating function can be captured at the level of the BOLD signal.^48^

Careful future experimentation is required to determine how memory plasticity changes as a function of task demands, contexts, goals, and individual differences. Extensions of the model to offline replay processes can serve to more precisely quantify the range and sequence of replayed events and sampling across brain regions during memory retrieval.

#### Other embeddings for memory graphs

The memory networks in the simulations were constructed by extracting semantic embeddings via word2vec from the Google News Corpus^11^. We initialized memory networks, graphs of items with weighted associations, based on the items used in the design of each human experiment. However, it is unclear whether word2vec is the most plausible proxy for a network of memory associations. For instance, the current embedding is bidirectional while real world associations can sometimes be asymmetrical. Furthermore, the word2vec method is suboptimal for capturing conditional or hierarchical relationships, such as those represented in prefrontal-hippocampal networks. Therefore, future research is required to implement and test more sophisticated embeddings, such as multi-scale successor representations^19^, Glove^49^, BERT^50^, ELMO^51^, or other bidirectional word embedding and neural NLP (Natural Language Processing) methods^52^.

#### Applications in computational psychiatry

SAMPL could provide a computational framework for understanding maladaptive memory function in psychiatric disorders, such as post traumatic stress disorder (PTSD)^53^. The model can be used to understand common symptoms of such disorders and design interventions. For instance, it may provide a process-based understanding of overgeneralization^54^ and flashbacks^55^. It may also help identify network properties underlying the over-accessibility of intrusive memories. For instance, this could be done by constructing a semantic embedding model and corresponding memory network from transcripts of a patient’s therapy sessions or diaries, focusing on the connections between neutral memories and intrusive target words. In conjunction with SAMPL, this information could help design computationally assisted interventions, such as targeted “forgetting attacks” designed to diminish the accessibility of memory subnetworks that are activated during undesirable intrusive thoughts. For instance, it would be possible to determine which series of memory items need to be activated in order to diminish the reachability, or alter the communicability distance^56^, of a particular memory node or subgraph within the memory network.

#### Summary

We present SAMPL, a simple model of memory using spreading activation and non-monotonic plasticity. SAMPL simulates the results of a series of human experiments on retrieval-induced forgetting, memory enhancement due to item and context repetition, and synchronization of memory across conversational networks with different structures. Importantly, we find that the suppressive dip of the non-monotonic plasticity function is required for capturing human behavioral findings. The model offers a simple but powerful framework, unifying seemingly contradictory accounts of memory.

## Acknowledgments

This project started at the MIND17 summer school at Dartmouth College, organized by Luke Chang and Jeremy Manning. We gratefully acknowledge Matteo Visconti di Oleggio Castello, Seth Koslov, and Anuya Patil, who contributed to and worked on related ideas during the summer school. This work was made possible thanks to funding by the John Templeton Foundation (IM), NIBIB R01EB022864, and NIMH Grant R01-MH104606 granted to Joshua Jacobs.

---

**Supplementary Figure 1.**
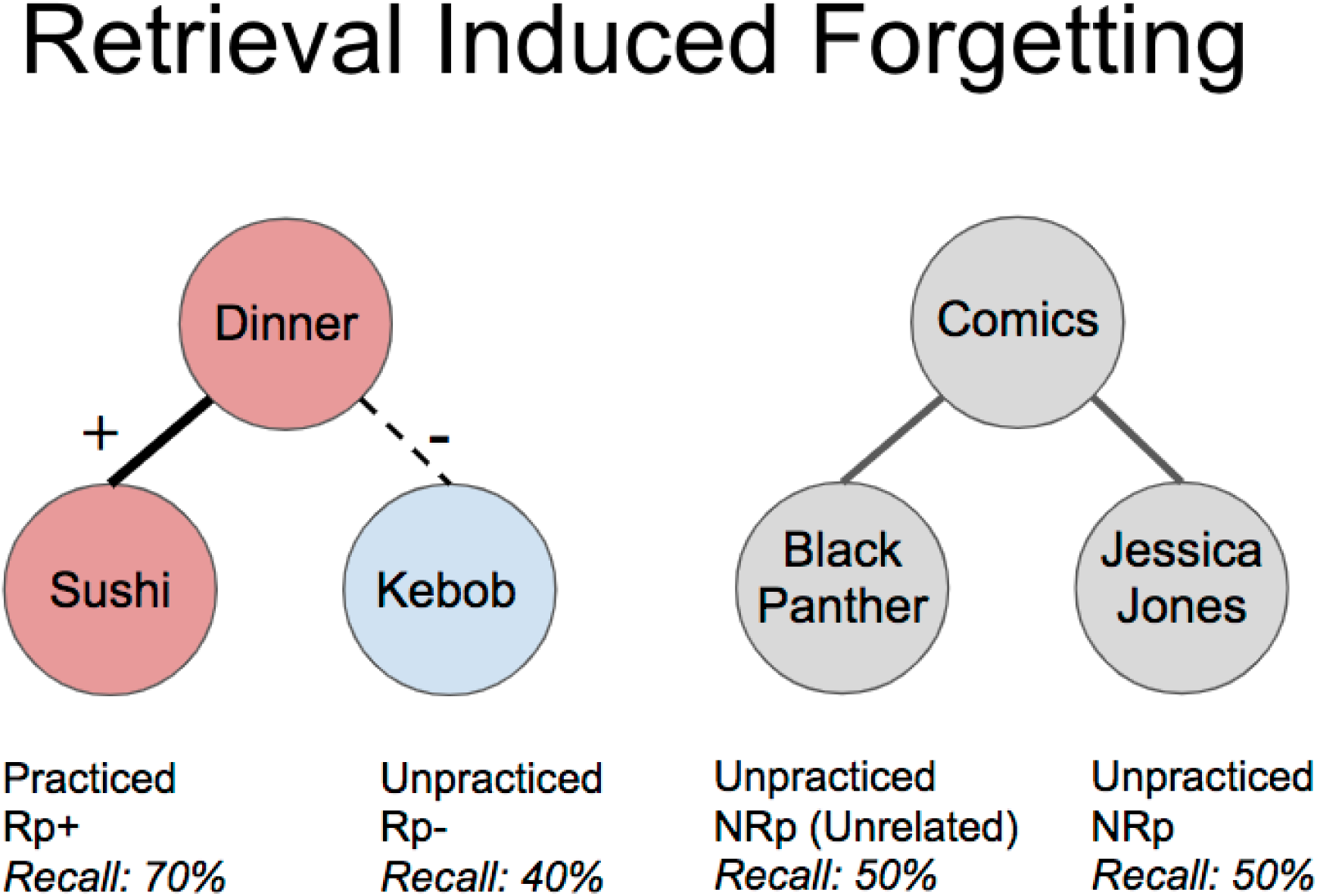
A schematic of a retrieval induced forgetting (RIF) experiment. Let us assume a simple memory network with two unconnected clusters: the dinner cluster with sushi and kebob as items, and the comics cluster, with black panther and jessica jones. Initially, let us assume that the retrieval for all items is equal, e.g., 50%. When sushi is activated, the strength of association between dinner and sushi increases. Therefore, sushi becomes memorable (from 50% to 70%). On the other hand, the weight of association with, and hence the memorability of, a related but unpracticed item, Kebob, drops (from 50% to 40%). Note that the comics cluster is not activated at all, and hence the memorability for Black Panther and Jessica Jones remain at 50%.

**Supplementary Figure 2.**
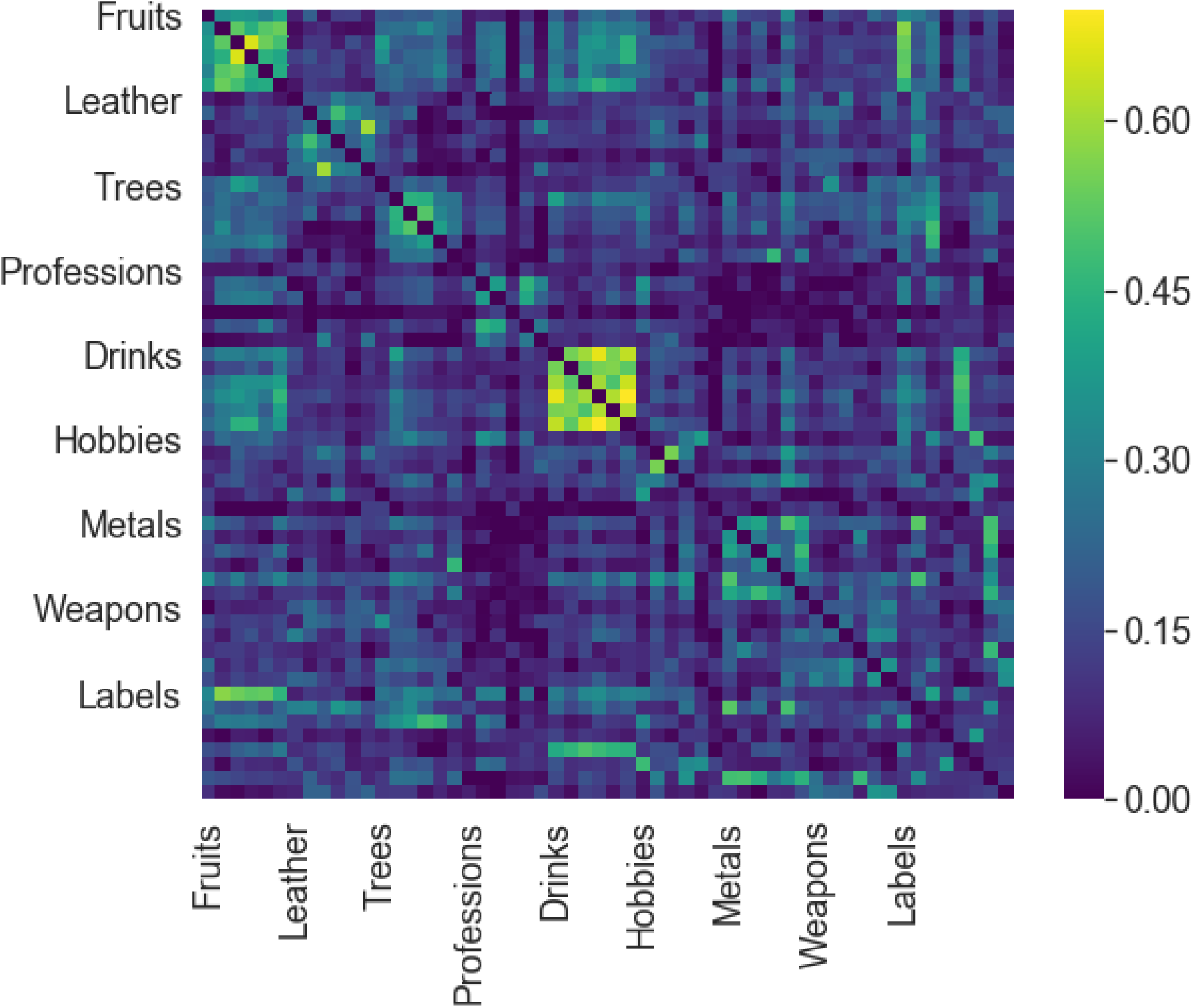
The initial state of the memory network for simulation. We constructed a memory network, displayed as a matrix of weights between pairs of items, based on the distances of the items in a word2vec embedding of Google news. Before the simulation, initial edge weights between every pair of items or exemplars were determined by word2vec cosine distances, and labeled by category. For a complete list of exemplars per category, see Supplementary Materials.

**Supplementary Figure 3.**
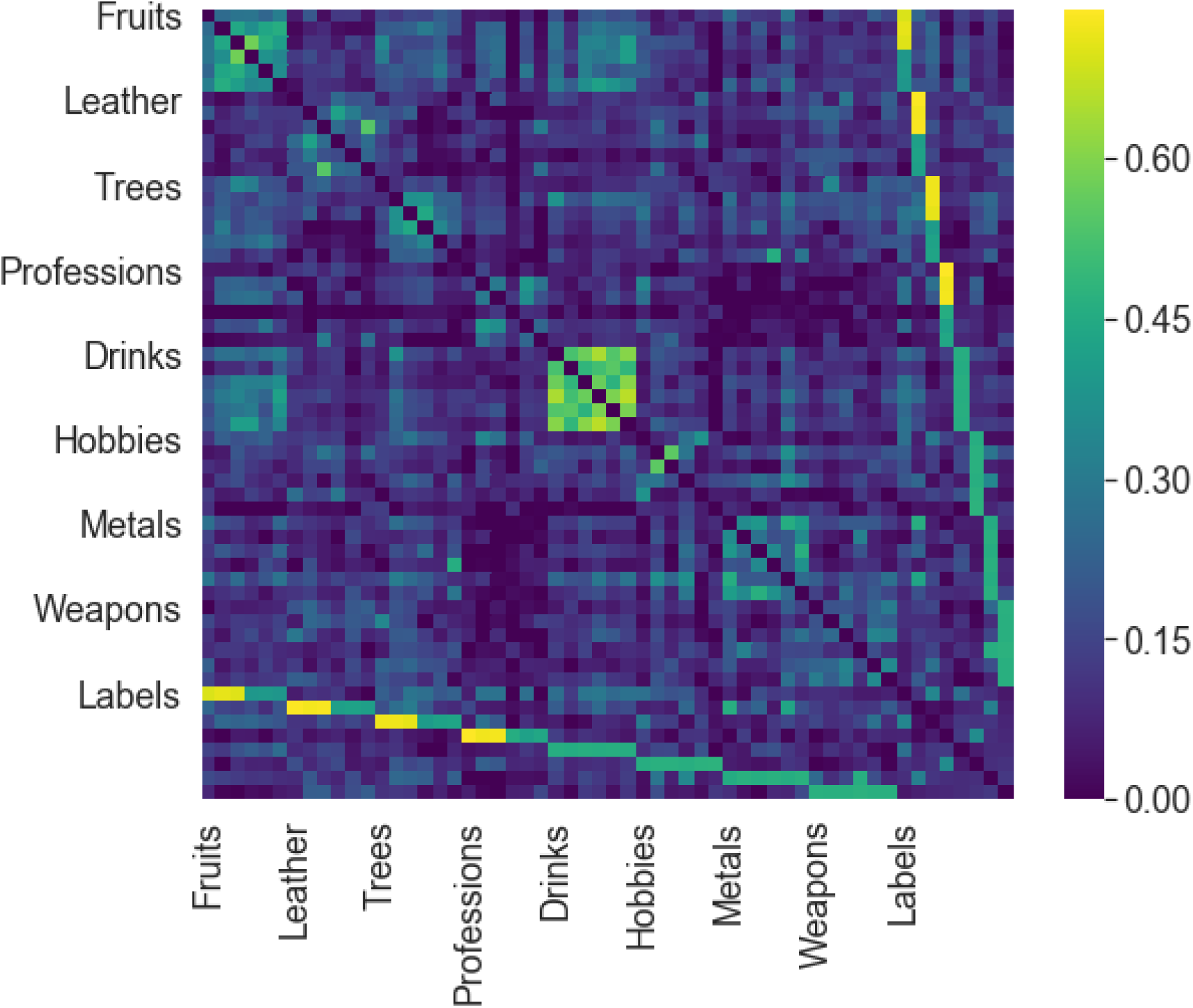
The final state of the memory network after simulation. Categorical relationships were maintained, and the largest weight changes were between items and their category labels.

**Supplementary Figure 3.**
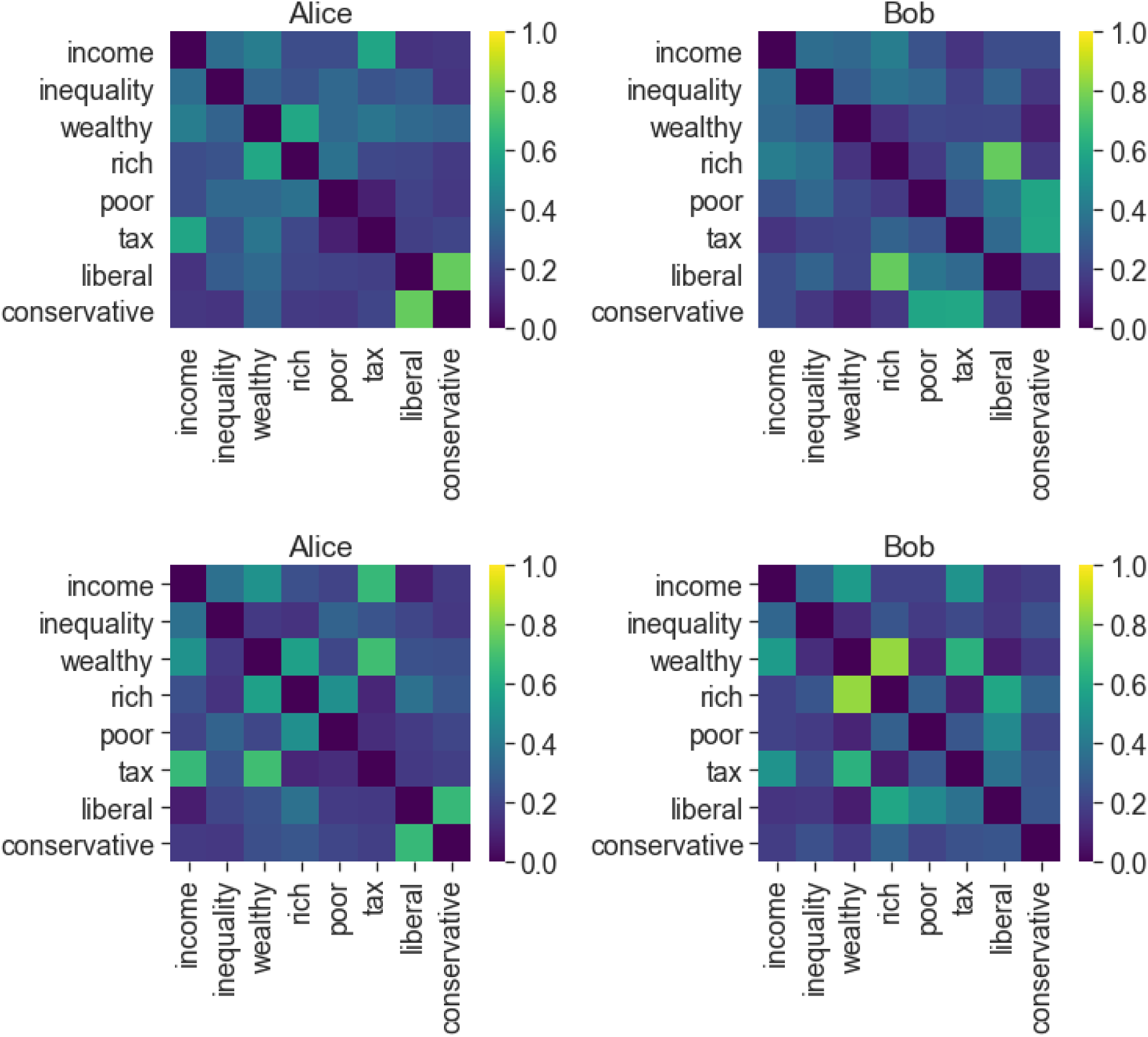
Memory network graphs before and after conversations. Typical edge weight matrices for both sender and receiver agents before conversation (top) and after conversation (bottom) are displayed. These simulation results are obtained when the sender agent selected words using the paired associate strategy.

**Supplementary Table 1.** Study 1 categories and exemplars

| Category | Exemplars |
| --- | --- |
| Fruit | orange, nectarine, pineapple, banana, cantaloupe, lemon |
| Leather | saddle, gloves, wallet, shoes, belt, purse |
| Tree | palm, hickory, willow, poplar, sequoia, ash |
| Profession | tailor, florist, farmer, critic, grocer, clerk |
| Drink | bourbon, scotch, tequila, brandy, gin, rum |
| Hobby | gardening, coins, stamps, ceramics, biking, drawing |
| Metal | chrome, platinum, magnesium, mercury, pewter, tungsten |
| Weapon | hammer, fist, lance, rock, arrow, dagger |

**Supplementary Table 2.** Grid search details

|  | Study 1, ABB (1994) | Study 2, SHS (2013) | Study 4, CMDG (2016) |
| --- | --- | --- | --- |
| **# simulations per parameter set** | 1 (no randomness) | 30 | 20 |
| **Suppression (y-minimum)** | -.75, -.5, -.4, -.3, -.2, -.1, -.05, -.025, -.01, 0 | 1.0, .8, .6, .5, .4, .3, .2, .1, 0 | 1.0, .8, .6, .5, .4, .3, .2, .1, 0 |
| **Enhancement (y-maximum)** | .75, .5, .4, .3, .2, .1, .05, .025, .01, 0 | -1.0, -.8, -.6, -.5, -.4, -.3, -.2, -.1, 0 | -1.0, -.8, -.6, -.5, -.4, -.3, -.2, -.1, 0 |
| **Dip center** | .05, .1, .15, .2, .25, .3, .35, .4 | .1, .2, .3, .4, .5 | .1, .2, .3, .4, .5 |
| **Dip width** | .05, .1, .15, .2, .25, .3, .35, .4 | .1, .2, .3, .4, .5 | .1, .2, .3, .4, .5 |
| **Discount ($\gamma$)** | .1, .2, .3, .4, .5, .75, .9 | .1, .2, .3, .4, .5, .7, .9 | .1, .2, .3, .4, .5, .7, .9 |
| **Learning rate** | N/A | N/A | .2, .4, .6, .8, 1.0 |

